# The draft genome sequence of *Eucalyptus polybractea* based on hybrid assembly with short- and long-reads reads

**DOI:** 10.1101/2021.05.18.444652

**Authors:** Teng Li, David Kainer, William J. Foley, Allen Rodrigo, Carsten Külheim

**Affiliations:** Research School of Biology, The Australian National University, Canberra 2600, Australia; Center for BioEnergy Innovation, Bioscience Division, Oak Ridge National Laboratories, Oak Ridge TN 37831 USA; College of Forest Resources and Environmental Science, Michigan Technological University, Houghton, 49931 MI USA

**Keywords:** Hybrid genome assembly, Eucalyptus, eucalyptus oil, nanopore long reads, genome

## Abstract

*Eucalyptus polybractea* is a small, multi-stemmed tree, which is widely cultivated in Australia for the production of Eucalyptus oil. We report the hybrid assembly of the *E. polybractea* genome utilizing both short- and long-read technology. We generated 44 Gb of Illumina HiSeq short reads and 8 Gb of Nanopore long reads, representing approximately 83× and 15× genome coverage, respectively. The hybrid-assembled genome, after polishing, contained 24,864 scaffolds with an accumulated length of 523 Mb (N50 = 40.3 kb; BUSCO-calculated genome completeness of 94.3%). The genome contained 35,385 predicted protein-coding genes detected by combining homology-based and de novo approaches. We have provided the first assembled genome based on hybrid sequences from the highly diverse *Eucalyptus* subgenus *Symphyomyrtus*, and revealed the value of including long-reads from Nanopore technology for enhancing the contiguity of the assembled genome, as well as for improving its completeness. We anticipate that the *E. polybractea* genome will be an invaluable resource supporting a range of studies in genetics, population genomics and evolution of related species in *Eucalyptus*.

## Introduction

Eucalypts (*Eucalyptus, Angophora* and *Corymbia* spp.) are economically important as the world’s most widely planted hardwood genera (Myburg et al. 2014). Globally, eucalypts are planted for pulp and paper production, timber, firewood and 1,8-cineole-rich essential oils (“Eucalyptus oil”), which are extracted from their leaves (Bauhus et al. 2010). They are also the dominant forest trees in Australia and thus of great ecological value. These genera are highly diverse with more than 800 species (Thornhill et al. 2019), ranging from the tallest flowering plant in the world (*E. regnans* over 100 m tall) to small shrubs (e.g. *E. vernicosa, E. yaltensis* and *E. surgens* < 2 m). Eucalypts have adapted to a wide range of climates in Australia and nearby islands from tropical to arid and temperate zones. They thrive in tropical monsoon (Köppen-Geiger classification Am), tropical rainforest (Af) and tropical savannah (Aw) regions with species such as *E. melanophloia, E. tereticornis* and *E. tetradonta*, respectively. Eucalypts are common in the arid zones which dominate Australia’s interior, with species such as *E. camaldulensis* (Bustos-Segura et al. 2017) surviving in the hot desert climate (BWh). In contrast, *Eucalyptus pauciflora* grows from coastal New South Wales and Victoria (Cfb) to the treeline of the Australian Alps (Wang et al. 2020; Woldendorp et al. 2008), which have a subpolar oceanic climate (Cfc). While some species occupy very narrow geographic and climate space (e.g. *E. polybractea*), others, such as *E. camaldulensis* (Bustos-Segura et al. 2017) and *E. tereticornis* (Bourne et al. 2017) cover vastly different climate space across the Australian continent.

Since the sequencing and annotation of the first *Eucalyptus genome, E. grandis* (Myburg et al. 2014), the focus of genetic and genomic research has been biomass accumulation (Cappa et al. 2013; Müller et al. 2019; Resende et al. 2012), wood properties (Cappa et al. 2013; Resende et al. 2017) and pest and pathogen resistance (Mangwanda et al. 2016; Oates et al. 2015; Resende et al. 2017), which are all important traits for the pulp and paper industry. Several studies have investigated species that are exclusively grown for the production of Eucalyptus oil by using genomic information based either on candidate genes in the terpene biosynthetic pathway (Padovan et al. 2017) or utilizing the related *E. grandis* genome sequence for reference mapping of reads (Kainer et al. 2019; Kainer et al. 2018). Due to high levels of synteny across the genus (Hudson et al. 2012), as well as shared ancestral genomic variants (Külheim et al. 2009), mapping of reads obtained from one species to another has been possible. However, regions that are unique to the non-reference species will not align, or align erroneously, and the genetic variants within those regions go undetected. Mapping quality has also improved with several iterations of editing fixed alleles in two eucalypt studies (Kainer et al. 2019; Orr et al. 2020).

Sequencing, assembly and annotation of *E. polybractea* will allow better short-read mapping of this and other closely-related species that are used for the production of essential oils. Having genome sequences from multiple species of eucalypts will also allow comparative studies, including variation of form (tall trees versus shrub), adaptation to climate, or the quantitative and qualitative variation of essential oils which is a well described trait in this genus (Padovan et al. 2014). Here we report the hybrid assembly of the genome of *E. polybractea* with high-coverage Illumina short reads (~ 83×) and low-coverage (~15×) Nanopore long reads using the MaSuRCA assembler. A schematic workflow of this study is shown in Fig. S1. We anticipate that the assembled and annotated genome of *E. polybractea* will be a catalyst for a range of genetics, population genomics, and evolution studies of this highly diverse genus and other related species.

## Material and Methods

### Sampling, Library Construction, and Sequencing

Leaf samples were collected from a *Eucalyptus polybractea* R.T. Baker progeny trial on the property of GR Davis Pty Ltd in West Wyalong NSW, Australia (33°58’S, 147°03’E). For Illumina short-read sequencing, leaves from 12 related (half-sib) individuals were flash frozen in liquid nitrogen and stored in −80°C until DNA extraction using the Qiagen DNeasy Plant kit (Qiagen, Valencia, CA, USA) as described in (Kainer et al. 2018). Short-read sequencing libraries were prepared using a modified protocol based on Rohland and Reich (2012) and were sequenced with 125 bp paired-end reads on the Illumina HiSeq 2500 platform at Macrogen (Republic of Korea). A total of 43.4 Gb of paired-end reads were generated from HiSeq data.

For long-read sequencing, branches were cut and placed in sealed bags for transport to Canberra, Australia, where they were placed in water at 4°C. Leaf sections were placed in a 2 ml Eppendorf tube with 5 mm steel beads, frozen in liquid nitrogen and ground in a Qiagen bead mill. DNA was extracted following the protocol of (Schalamun et al. 2018). DNA was quantified using a Qubit fluorimeter and quality was assessed by separation of the DNA on a 0.7% agarose gel. Due to high foliar concentrations of plant secondary metabolites, particularly tannins and essential oils, purification steps by either SPRI beads or precipitation with chloroform/isoamylalcohol were repeated where necessary. Oxford Nanopore long reads were generated by using the Ligation Sequencing 1D Kit (SQK-LSK108) with approximately 5 μg high quality DNA, which was extracted from four individuals due to varying success of high-quality native DNA extraction. Initial efforts to extract native DNA from the same individuals as for short-read sequencing failed, likely due to high foliar concentrations of eucalyptus oil. We then selected individuals that had the lowest concentration of foliar eucalyptus oil (Kainer et al. 2019). Four libraries were prepared and sequenced on four different flowcells using the MinION device (Oxford Nanopore, Oxford, UK), Nanopore reads were basecalled from their raw FAST5 files using Albacore v2.3.1. Applying a minimum length cutoff of 500 bp and discarding reads with quality <7, this study produced a total of 7.96 Gbp (N50: 16.1 kb, longest read: 145,007 bp).

### Genome Characteristics

Jellyfish v. 2.2.6 (Marcais and Kingsford 2011) was employed to generate the frequency distribution of 17-, 21-, and 25-mers of the clean short reads obtained from paired-end Illumina sequencing libraries, and these distributions were subsequently processed by GenomeScope (Vurture et al. 2017). The analysis shows that the genome size of *Eucalyptus polybractea* is between 490 and 543 Mb with a population heterozygosity level of 2% (Fig. S2). The genome sizes estimated by kmer-based statistical approach are within the range of sizes listed for other *Eucalyptus* species (381 Mb - 724 Mb) as reported on the Plant C-values Database (http://data.kew.org/cvalues).

### Genome assembly

*Eucalyptus polybractea* genome was assembled *de novo* using Maryland Super-Read Celera Assembler v.3.2.6 (MaSuRCA) (Zimin et al. 2013) with short reads only, followed by a hybrid (Illumina + Nanopore data) assembly method. The short reads used for both assemblies derived from paired-end libraries which were trimmed for adapters by BBDuk(BBMap), but not trimmed for any aspects of quality, cleaning or error correction. The assembler MaSuRCA constructs long and accurate mega-reads from the combination of long and short read data, essentially converting each Nanopore long read into one or more very long, highly accurate reads (Zimin et al. 2017). The short-reads-only and hybrid assemblies yielded total assembly length of 1,063 Mb and 523 Mb, respectively (Table 1). The guanylic and cytidylic acid (GC) content of the *E. polybractea* hybrid assembly genome (mean = 39.0%) is similar to the *E. grandis* genome (39.3%). The hybrid assembly was subsequently polished by Pilon v.1.22 (Walker et al. 2014), which uses information in short reads to correct bases, fix mis-assemblies, and fill gaps. Pilon was applied for multiple iterations until the polished genome remained unchanged. Compared to the short read only assembly, the inclusion of approximately 15× genome coverage of Nanopore long reads for a hybrid assembly led to a 95% decrease in the number of scaffolds from 502,024 to 24,864 scaffolds and an 11-fold increase in the scaffold N50 length from 3,393 bp to 40,281 bp (Table 1).

**Table 1:** Summary of genome assembly and annotation of *Eucalyptus polybractea*.

|  | Illumina + Nanopore | Illumina only |
| --- | --- | --- |
| Genome assembly |  |  |
| Number of contigs | 24 864 | 502 024 |
| Total length | 523 113 982 bp | 1063 617 135 bp |
| N50 size | 40 281 bp | 3393 bp |
| Longest | 580 368 bp | 73 252 bp |
| GC (%) | 39.0 | 38.9 |
| BUSCO genome completeness |  |  |
| Complete | 2001 (94.3%) | 1426 (67.2%) |
| Complete and single copy | 1765 (83.2%) | 1177 (55.5%) |
| Complete and duplicated | 236 (11.1%) | 249 (11.7%) |
| Fragmented | 21 (1.0%) | 408 (19.2%) |
| Missing | 99 (4.7%) | 287 (13.6%) |
| Genome annotation |  |  |
| Number of protein-coding genes | 35 385 |  |
| Mean transcript length | 2878.6 bp |  |
| Mean coding sequence length | 1373.3 bp |  |
| Mean exon length | 313.6 bp |  |
| Mean intron length | 445.5 bp |  |

### Repeat content analysis

Repetitive sequences in the *E. polybractea* genome were identified and classified using a combination of *de novo* and homology-based approaches. Initially, a *de novo* repeat library of *E. polybractea* was built by RepeatModeler v. 1.0.4 (Smit and Hubley 2008-2015) with default parameters based on the hybrid assembly. RepeatMasker v. open-4.0.7 (Smith et al. 2013-2015) was then employed to detect repeats in the *E. polybractea* genome, by aligning sequences from the hybrid assembly to the RepeatMasker combined database (Dfam_consensus (Hubley et al. 2016) and Repbase (Jurka et al. 2005)) as well as to the *de novo* repeat library that we built. Repeat sequences were estimated to account for 40.1% (210 Mb) of the *E. polybractea* genome (Table S1), with long terminal repeats (LTRs) being the most pervasive class (17.6%). However, DNA transposons encompass only 4.1% of the genome. In addition, the percentage of repetitive elements in *E. polybractea* is less than that in *E. grandis* (50.1%), which likely accounts for the smaller genome size of *E. polybractea* (523 Mb) compared to *E. grandis* (640 Mb).

### Genome annotation

The gene annotation for the *E. polybractea* genome was performed by combining homology-based and *de novo* approaches. For the homology-based method, the hybrid assembly of the genome was aligned to the protein-coding genes predicted in *E. grandis* (Myburg et al. 2014) using TBLASTN v. 2.2.31 (Camacho et al. 2009). The homologous genome sequences were then aligned against the matching protein sequences by GeneWise v. 2.4.1 (Birney et al. 2004) to obtain accurate spliced alignments. The *de novo* prediction was estimated by the *ab initio* gene predictors Augustus v. 3.2.1 (Stanke et al. 2006) and GenScan (Burge and Karlin 1997), followed by filtering partial sequences and genes with length less than 100 bp. The final consensus protein-coding gene set of the *E. polybractea* genome was generated by EVidenceModeler (EVM) v. 1.1.1 (Haas et al. 2008), with the combined gene annotation results of *de novo* and homology-based predictions. Our results showed that 35,385 protein-coding genes were identified in the *E. polybractea* genome, with an average coding length of 1,373 bp (Table 2).

**Table 2.** Genome annotation of the hybrid assembly *E. polybractea* genome.

| Method | Gene<br>number | Mean<br>transcript<br>length (bp) | Mean<br>CDS<br>length (bp) | Mean<br>exon<br>per gene | Mean<br>exon<br>length (bp) | Mean<br>intron<br>length (bp) |
| --- | --- | --- | --- | --- | --- | --- |
| **Homology to *E. grandis*** | 35,490 | 2702.3 | 1330.4 | 4.2 | 313.2 | 422.4 |
| ***De novo* by AUGUSTUS** | 28,157 | 2888.5 | 1231.1 | 5.0 | 244.4 | 410.5 |
| ***De novo* by GENESCAN** | 26,098 | 2774.7 | 1455.4 | 3.4 | 423.0 | 540.6 |
| **EVidenceModeler (EVM)** | **35,385** | **2878.6** | **1373.3** | **4.4** | **313.6** | **445.5** |

## Results and Discussion

### Assessment of the genome assembly

To evaluate the completeness of the genome assembly, Benchmarking Universal Single-Copy Orthologs v.3.0.2 (BUSCO) (Simao et al. 2015) was employed to search against the Eudicotyledons database of 2,121 orthologs. The results showed that the genome completeness was substantially improved in the hybrid assembly compared to the short reads only assembly, from 67.2% to 94.3% of the complete sequences detected by BUSCO. In addition, 1,765 single-copy and 236 duplicated orthologs were observed in the hybrid assembly with only 21 (1%) fragmented genes (Table 1).

In order to validate our genome assembly by the hybrid approach, comparison was performed between the *E. polybractea* genome and *E. grandis* genome. The protein-coding genes predicted in *E. grandis* genome (Myburg et al. 2014) were mapped to the hybrid assembly of *E. polybractea* by BLAT algorithm with default parameters (Kent 2002). Statistical analysis was performed at different levels of percentage of sequence homology and percentage of coverage, which demonstrated that the hybrid assembly of *E. polybractea* covered approximately 98% of the protein-coding genes in *E. grandis* (Table S2). Further, we mapped the clean short reads from Illumina HiSeq sequencing of 12 related (half-sib) individuals to the hybrid assembly of the *E. polybractea* genome using the Burrows-Wheeler Aligner v0.7.17-r1188 (BWA) (Li 2013), and extracted the mapping rate and coverage rate from the result by Qualimap v2.2.1 (Okonechnikov et al. 2016). The results showed that our hybrid assembly had high mapping rate of the short reads (97.5%) with a mean depth of 49.3x, while 91.6% of the assembled genome had a mean depth over 10x (Fig. S3), ensuring a high level of mapping accuracy at the nucleotide level. Additionally, we also performed variant calling with the BWA mappings of the 12 half-sib individuals using the Genome Analysis Toolkit v3.5 (DePristo et al. 2011), resulting in the detection of 12,983,296 variants. The mapping was followed by filtering of adjacent indels within 5 bp and low quality (quality score less than 30, sequencing depth either less than one third of mean depth or three times bigger than the mean depth) single nucleotide variants (SNVs) using BCFtools (Li 2011), resulting in the detection of 10,740,778 SNVs that passed filtering. The results revealed that the population heterozygosity level of the *E. polybractea* genome was approximately 2.05%. In summary, all of these analyses indicated that our draft genome sequence has high contiguity, accuracy, and, more importantly, a high degree of gene space completeness.

## Data availability

Raw Illumina and Nanopore reads sequenced in this study are available in the Short Read Archive, together with the Whole Genome Shotgun project have been deposited at NCBI under BioProject PRJNA525896. The assembled genome is available at GenBank under the accession number SMSX00000000. All supplementary figures and tables are provided in additional file 1.

## Acknowledgements

The authors would like to acknowledge Ms Miriam Schalamun and Dr Benjamin Schwessinger for aiding in the development of the native DNA extraction protocol.

## Declarations

### Consent for publication

All authors have read and approved the manuscript for submission to Tree Genetics and Genomes.

### Code availability

All tools used in this study were properly described in the Methods section together with the versions used.

### Authors’ contributions

The study was conceived by CK, WJF and AR. DNA for short-read sequencing was extracted by DK, DNA for long-read sequencing was extracted by CK and sequenced by CK. TL did the bioinformatics analyses of this study and wrote the draft of the MS with DK and CK. All authors read and approved the final version of this manuscript.

