## Supplemental Material for "The draft genome sequence of *Eucalyptus polybractea* based on hybrid assembly with short- and long-reads reads"

### Title

### Affiliations

Table S1. Classification of repetitive elements in the hybrid assembly of *E. polybractea* genome.

| Type | Combined[a] | | *De novo* | | RepeatMasker (Dfam_consensus & Repbase) | |
|---|---|---|---|---|---|---|
| | **Repeat length (bp)** | **% of genome** | **Repeat length (bp)** | **% of genome** | **Repeat length (bp)** | **% of genome** |
| **SINEs** | 2,853,743 | 0.55 | 2,826,159 | 0.54 | 27,584 | 0.01 |
| **LINEs** | 12,666,063 | 2.42 | 12,661,936 | 2.42 | 9,295,794 | 1.78 |
| **LTR** | 91,928,860 | 17.58 | 91,794,854 | 17.55 | 80,599,946 | 15.41 |
| **DNA transposons** | 21,627,690 | 4.14 | 21,444,864 | 4.10 | 10,515,214 | 2.01 |
| **Unclassified** | 80,431,369 | 15.38 | 80,341,410 | 15.36 | 1,143,082 | 0.22 |
| **Total[a]** | 209,507,725 | 40.06 | 209,069,223 | 39.98 | 101,581,620 | 19.43 |

[a]Total repeats contained all the repeats been detected. As some overlaps existed between different methods, the total repeats may be shorter than the sum of repeats separately detected.

Table S2. Evaluation of gene space completeness for the *Eucalyptus polybractea* genome, with aligned to the protein-coding genes predicted in *E. grand*is genome.

| Gene length | Total number | Total Length (bp) | Covered by assembly | With >90% sequence in one scaffold | With >50% sequence in one scaffold |
|---|---|---|---|---|---|
| >0 | 46,280 | 54,521,616 | 97.4% | 42,766 (92.4%) | 44,983 (97.2%) |
| >200 | 46,280 | 54,521,616 | 97.4% | 42,766 (92.4%) | 44,983 (97.2%) |
| >500 | 36,373 | 51,248,055 | 97.6% | 34,070 (93.7%) | 35,524 (97.7%) |
| >1,000 | 23,124 | 41,295,171 | 97.8% | 21,827 (94.4%) | 22,642 (97.9%) |

Figure S1. The genome assembly and annotation pipeline of *Eucalyptus polybractea*.

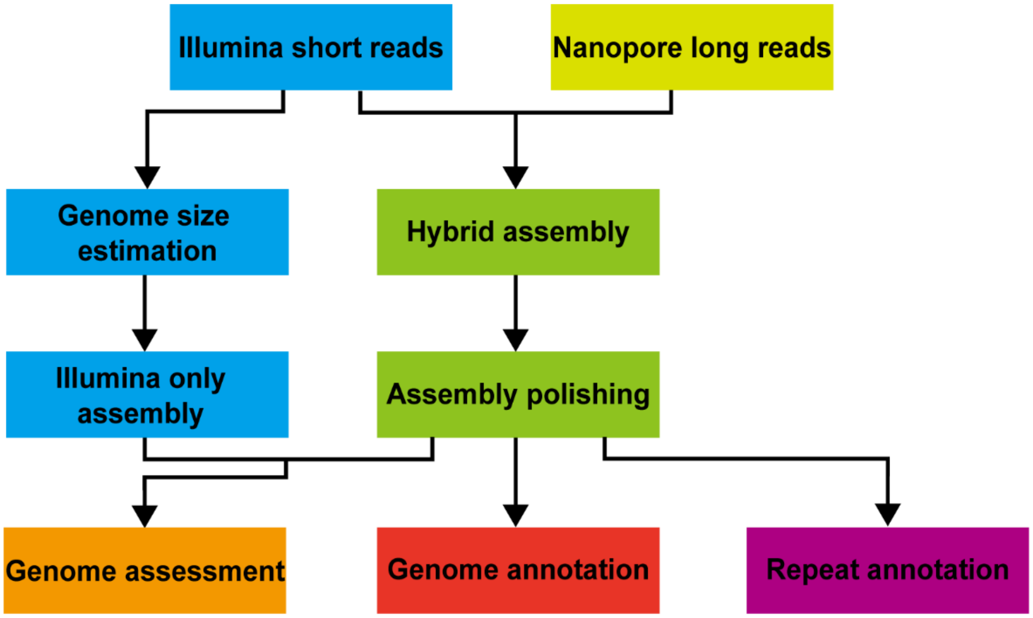

Figure S2. Estimate the genome size, repeat content, and heterozygosity of *E. polybractea* using GenomeScope based on 17-mer (A), 21-mer (B), and 25-mer (C) in HiSeq sequence reads.

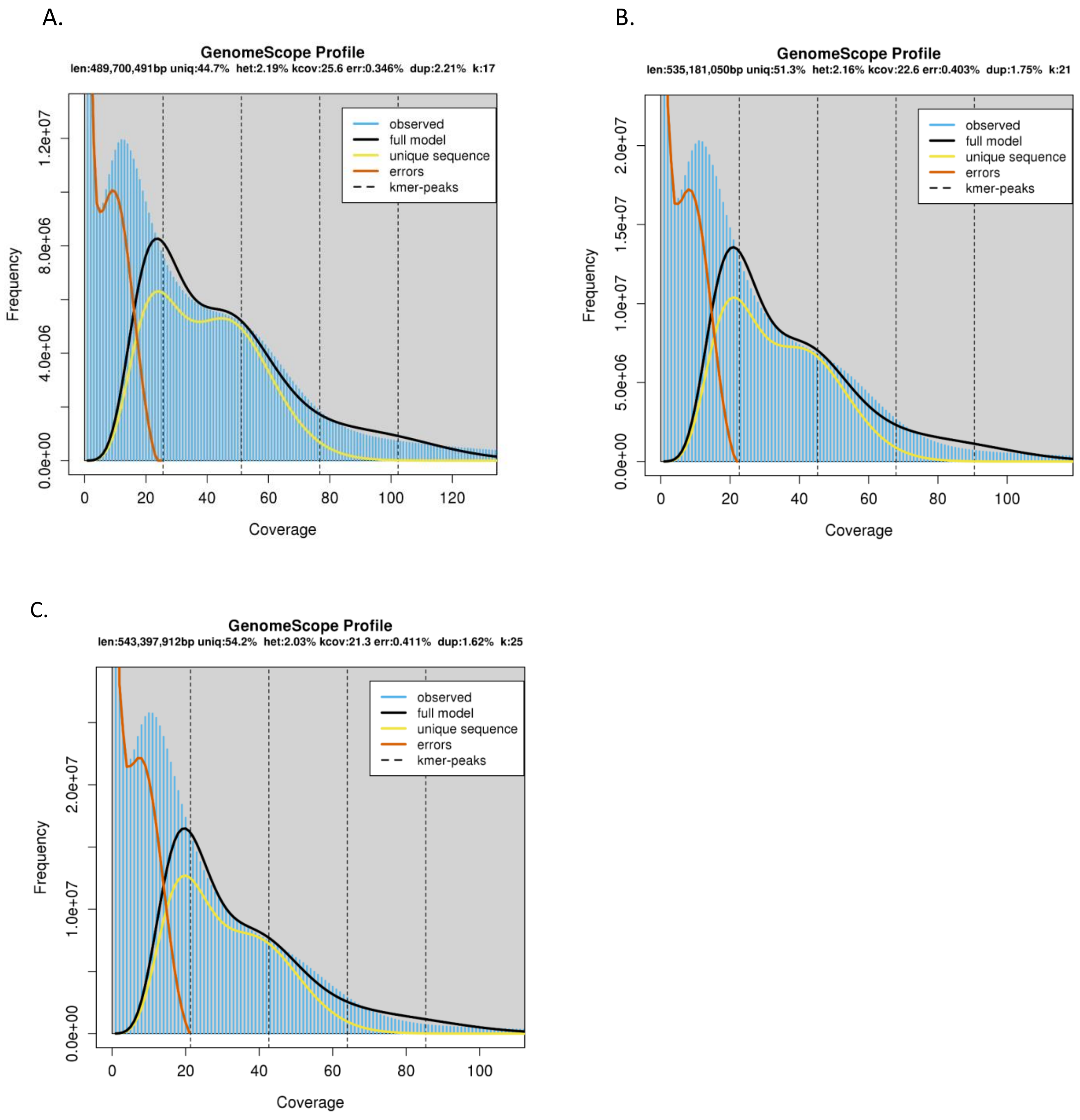

Figure S3. Sequencing depth distribution for the *E. polybractea* genome. The HiSeq sequence reads were realigned onto the hybrid assembly using the BWA software. The sequencing depth of each base was calculated and plotted.

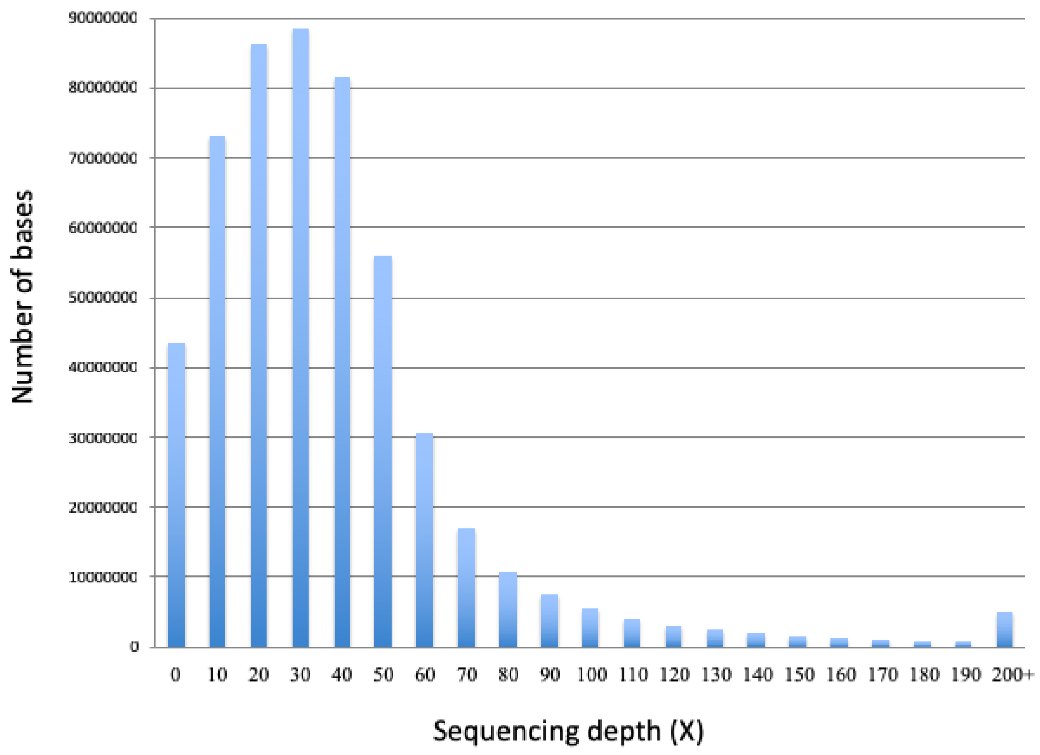
